# Probiotic consumption relieved human stress and anxiety symptoms via modulating the gut microbiota and neuroactive potential

**DOI:** 10.1101/2020.08.05.237776

**Authors:** Teng Ma, Hao Jin, Lai-Yu Kwok, Zhihong Sun, Min-Tze Liong, Heping Zhang

**Author notes:** Correspondence: Heping Zhang.

## Abstract

Stress has been shown to disturb the balance of human intestinal microbiota and subsequently cause mental health problems, such as anxiety and depression. The gut microbial communities are able to synthesize and/or consume various neuroactive metabolites, and preliminary human studies have also demonstrated the ability of probiotics to modulate the levels of neurotransmitter. However, the study and interpretation of the biological significance of microbial neuroactive compounds have been hindered by the lack of dedicated reference databases and corresponding human intestinal microbiota reference genomes. Our previous study showed that ingesting the probiotic strain, *Lactobacillus* (*L*.) *plantarum* P-8, for 12 weeks could alleviate stress and anxiety of stressed adults. The current study was a follow-up work aiming to further elucidate mechanisms behind the observed beneficial effects by performing deep analysis of the fecal metagenomes of the probiotic (n = 43) and placebo (n = 36) groups. Comparing with the probiotic group, the gut microbiomes of the placebo group showed significantly higher Bray-Curtis dissimilarity between weeks 0 and 12 (*p* < 0.05). Comparing with week 0, the Shannon diversity index of the placebo group decreased significantly at week 12 (t-test; *p* < 0.05), but such decrease was non-significant for the probiotic group. Additionally, the fecal metagenomes of the probiotic group showed significant increases in the species-level genome bins (SGBs) of *Bifidobacterium adolescent*, *Bifidobacterium longum*, and *Fecalibacterium prausnitzii* increased, while decreases in the SGBs of *Roseburia faeci* and *Fusicatenibacter saccharivorans*. Furthermore, the 12-week probiotic supplementation enhanced the diversity of neurotransmitter-synthesizing and/or -consuming SGBs, and the levels of some predicted microbial neuroactive metabolites (short chain fatty acids, gamma-aminobutyric acid, arachidonic acid, and sphingomyelin et.al). In conclusion, this study revealed the potential mechanism of probiotics in alleviating stress and anxiety via the gut-brain axis. The modulation of the intestinal microbiota by probiotics is an attractive strategy for managing stress and anxiety.

## Introduction

Stress is a ubiquitous part of human daily life that can be defined as the disruption of homeostasis due to environmental factors, physical or psychological stimuli. It can cause anxiety and even precipitate depression in serious cases[1, 2]. Depression affects over 300 million people among the global population, accounting for almost 800,000 suicidal deaths per year[3]. Mental illnesses impose an enormous health care costs and high burden of disease. Recent studies have associated mental illness risk with the commensal microorganisms residing in the gastrointestinal tract or the gut microbiota[4, 5]. The gut microbiota can be viewed as a reservoir containing a collective genome that harbors at least 100 times as many genes as the human genome[6]. The co-evolution of the human genome and the gut microbiome has led to the establishment of complex bidirectional interactions between the gastrointestinal tract, enteric nervous system, and central nervous system coordinated by different organ systems[1, 7]. Interestingly, the intestinal microbiota widely participates in the synthesis and release of various hormones and gut-brain axis-related neurotransmitters, which in turn modulate the brain function and host behavior[5, 8]. Gastrointestinal disturbances have also been proven to be related to mental illnesses, including stress, anxiety, depression, Parkinson’s and Alzheimer’ diseases (P/AD)[9].

Probiotics are defined as living microorganisms that produce health benefits to the host when administered in adequate amounts[10]. Since probiotics have been shown to modulate the host gut microbial communities and their synthesis/release of certain neuroactive compounds, probiotic-based therapy has thus been proposed as an alternative treatment for some neurological and neurodegenerative diseases[11, 12]. Recently, the administration of some probiotic strains has shown beneficial effects in relieving stress, anxiety, and depression symptoms[13, 14], particularly, strains belonging to the *Bifidobacterium* (*B*.) and *Lactobacillus* (*L*.) genera, which have long history of safe use and proven health-promoting functions[15]. However, most of these studies have focused on exploring their impact on host phenotypes by administering behavior tests, cognitive assessments and questionnaires to assess relief of clinical symptoms[14, 16]. Some studies have attempted to characterize probiotic-induced changes in the taxonomic (16S rRNA-based) profiles of intestinal microbiota[17, 18], but failed to provide a comprehensive overview of the gut microbiota function and neuroactive potential due to the lack of corresponding reference genomes[19]. Microbial reference genomes are indispensable tools for estimating the abundance of certain microbes in the metagenomes and deciphering their ecological role and function in the microbiome[20]. Recent studies have successfully reconstructed metagenome-assembled genomes (MAGs) from metagenomes based on the binning theory, which has massively expanded the catalogue of human gut reference genomes[21, 22]. The availability of a comprehensive human gut genome catalogue is critical in unveiling the impact of gut microbiome in human health and disease through population and functional analysis[23].

Previous animal studies or studies based on correlative analysis in patients have undoubtedly shown that the gut microbiota plays an influential role in many stress-related conditions; however, the causal relationship and extent of contribution of gut microbiome in stress-related medical conditions still need to be established by further human clinical trials[24]. Previously, our research team performed a 12-week randomized, double-blind, and placebo-controlled human trial demonstrating the clinical efficacy of the probiotic strain, *L. plantarum* P-8, in ameliorating stress- and anxiety-related symptoms in stressed adults[25]. Moreover, comparing with the placebo group, the 12-week treatment of P-8 resulted in a marginal decrease in plasma cortisol but significantly higher magnitudes of reduction in pro-inflammatory cytokines, such as IFN-γ and TNF-α. These physiological changes were accompanied by enhanced memory and cognitive traits, such as social emotional cognition and verbal learning and memory, upon administration of P-8. However, the mechanism behind the clinical symptom improvement remained unknown. The current study was a follow-up work that aimed to elucidate such mechanism from the perspective of the regulation of intestinal microbiota. This study based on in-depth analysis of the gut metagenomes of subjects participating in the clinical trial. Data obtained from this work would also reveal the potential functionary mechanism of probiotics in interacting with the host in relieving stress and anxiety.

## Materials and Methods

### Experimental design and subject recruitment

The current study was a follow-up work of our previous randomized, double-blind, and placebo-controlled human trial[25]. Briefly, our previous study assessed a total of 201 subjects for their eligibility of trial. Inclusion criteria: men or women, aged 18-60 years old, body mass index within a healthy range, no severe illnesses, willing to comply and follow throughout the study, and perceived moderate level of stress based on PSS-10. Exclusion criteria: individuals suffering from type-I diabetes, severe illnesses, HIV/AIDS, and glucose-6-phosphate dehydrogenase deficiency; and subjects who were not likely to complete the study based on the investigators’ opinion. Sixty-nine subjects were excluded after screening according to the inclusion and exclusion criteria. The remaining 132 subjects were randomized into two groups [probiotic (P-8) and placebo; n=66 per group]. During the trial period, 14 and 15 subjects were excluded from the study due to reasons like voluntary dropout, non-compliance in answering questionnaires, providing blood samples and/or completing the computerized CogState Brief Battery (CBB) test. 52 probiotic-receivers and 51 placebo-receivers finished the 12-week clinical trial. However, 9 probiotic-receivers and 14 placebo-receivers did not provide a fecal sample at week 12. These subjects were then excluded from the current analysis, and the final cohort included 43 probiotic-receivers and 36 placebo-receivers (Table S1).

The intervention was a daily dose of two grams of *L. plantarum* P-8 (2×10^10^ CFU/sachet/day; malodextrin as excipient) or placebo material (only malodextrin; light yellow powder with identical taste and appearance as the probiotic material); each dose was packaged in an individual aluminium sachet, stored away from direct sunlight and below 30°C, and the treatment lasted 12-weeks. Both the probiotic and placebo materials were manufactured by JinHua YinHe Biological Technology Co. Ltd., China under ISO9001 and HALAL standards. HALAL certification was obtained from ARA HALAL Development Services Center Inc. (ARA), which was recognized by JAKIM, Malaysia.

### Extraction of DNA, metagenomic sequencing and quality control

Metagenomic DNA was extracted from approximately 100 mg of stool using the QIAamp Fast DNA Stool Mini Kit (Qiagen GmbH, Hilden, Germany) following the manufacturer’s instruction, and the quality of DNA extraction was examined by 0.6% agarose gel electrophoresis and a Nanodrop spectrophotometer (ratio of absorbance at 260 nm/ 280 nm). Shotgun metagenomic sequencing was performed using an Illumina HiSeq 2500 instrument. Libraries were constructed by DNA fragments of approximately 300 bp length, and paired-end reads were generated using 100 bp in the forward and reverse directions. Low-quality reads, adaptor sequences, and host contaminating reads were removed.

### Metagenomic assembly, contig binning, and genome dereplication

All samples were processed with the standard quality control setting employed by metaSPAdes (v. 3.13.0)[26], with the options −k 33,55,77,99,111 -meta. To evaluate the results of metagenomic assembly, QUAST (v.5.0.0)[27] was used with the parameters, “--min-contig 2000”, and assembled scaffolds larger than 2,000 bp were selected for binning. Afterwards, MAGs were generated for each sample using two tools, MaxBin2 (v2.2.1)[28] and MetaBAT2 (v.2.12.1)[29], with default options by means of a combination of sequence composition and coverage information. Then, the MetaWRAP’s bin refinement module (v.1.1.8)[30] was used to combine and improve the results generated by the two binners. The raw reads were mapped back to the corresponding assemblies through BWA-MEM (v.0.7.17-r1188)[31], and the read depths were calculated using samtools (v.1.6; ‘samtools view -bS’ followed by ‘samtools sort’)[32] together with the jgi_summarize_bam_contig_depths function from MetaBAT2.

The levels of completeness and contamination of each MAG were evaluated with CheckM (v.1.0.722)[33] using the lineage_wf workflow. The quality of MAGs were classified as high (completeness≥80%, contamination≤5%), medium (completeness≥70%, contamination≤10%), and partial (completeness ≥ 50%, contamination ≤ 5%) based on the quality standards defined in Parks et al., 2017[34]. Genomic comparisons were performed by the dRep tool (v.2.2.2)[35] with options -pa 0.95 (primary cluster at 90%) -sa 0.95 (secondary cluster at 95%) to identify groups of essentially identical genomes and select the best genome from each replicate set. The results generated by dRep were used to extract species-level genomes bins (SGBs) from the high-quality genome dataset based on single representative genomes.

### Taxonomic annotation and phylogenetic analysis

Taxonomic classification and gene annotation were performed on the scaffolds of MAGs. Briefly, the scaffolds were annotated using the Kraken 2 tool (v.2.0.8-beta)[36] and NCBI nonredundant Nucleotide Sequence Database (NT, downloaded on 2019.04.21) with default parameters. The genes in scaffolds were predicted using Prodigal (v.2.6.3; meta option)[37]. Then, the predicted genes were searched against the UniProt Knowledgebase (UniProtKB, release 2019_04) using the blastp function of DIAMON[38].

The phylogeny analysis was performed based on the 400 universal PhyloPhlAn markers, applying the parameters described by Pasolli et.al[22]. The generated phylogenetic tree was visualized using iTOL[39].

### Calculation of abundance of SGBs

The BBMap tool (https://github.com/BioInfoTools/BBMap) was used to map raw reads to the scaffolds with the parameters “minid=0.95 idfilter=0.95”. The coverage of scaffolds was calculated by pileup.sh using sam file as input. The unit, reads per kilobase per million (RPKM), calculated by an in-house script was used to indicate the average content of each scaffold in each SGB.

### Identification of neuroactive compounds

A metabolic reconstruction was performed based on literature (gut-brain modules, GBMs)[19] and the metabolite database, MetaCyc[40], aiming to convert the genome information obtained from metagenomic binning into neuroactive potential and microbial metabolic function. For each SGB, open reading frames (ORFs) were predicted using Prodigal (v.2.6.3; meta option) with default parameters. Annotation of metabolic potential and pathways, e.g., pathways related to the metabolism of carbohydrate and amino acid, was based on key reactions in the Kyoto Encyclopedia of Genes and Genomes (KEGG) Orthologies (KOs) database (as of 2018). Each GBM corresponded to the process of synthesis or release of a neuroactive compound by specific gut microbes. All the pathways were detected by the Omixer-RPM tool (v.1.0)[41] and the pathways were defined in SGBs based on the parameter -c 0.66.

### Prediction of microbial metabolites

Microbial sequence features were used to predict the gut metabolomic profiles. Firstly, 1,000,000 reads from each sample were subsampled using the seqtk tool (v. 1.3-r106; https://github.com/lh3/seqtk), and the subsampled reads were compared using the blastx function of DIAMON with options ---query-cover 90 --id 50 (same as the parameter settings of HUMANN2 comparison process). Then, the best hit of each gene was selected, and the abundance of each gene in each sample was calculated. Finally, the MelonnPan-predict workflow (Model-based Genomically Informed High-Dimensional Predictor of Microbial Community Metabolic Profiles) was employed to convert microbial gene abundances into a predicted metabolomic table[42].

### Data and code availability

All sequencing data generated in this study have been deposited in NCBI-SRA (BioProject: PRJNA634666). The analysis codes have been made available under https://github.com/TengMa-Cleap/Probiotics-relieve-human-stress-and-anxiety-project.

## Results

### Metagenomic assembly strategy and general genomic characteristics

A total of 158 stool samples were shotgun sequenced, and the samples were collected at weeks 0 and 12 from each of the 79 subjects (n = 43 and 36 for P-8 and placebo groups, respectively; Table S1). This study generated a total of 1.52 Tbp high-quality paired-end reads (8.34 ± 1.41 Gbp per sample; range = 4.72 - 11.96 Gbp), The data of each sample were singly assembled into scaffolds using MetaSPAdes (average N50 length = 17.19 Kbp; Table S2). The metagenomic binning with MetaBAT2 and MaxBin2 yielded 14,534 and 10,545 raw bins from the initial scaffolds, respectively. Then, the bin_refinement module from MetaWRAP was employed to merge the compatible bins generated by the two binning tools, and 6566 bins were resulted. Among them, 3803, 851, and 1240 were assigned to high-, medium-, and partial-quality MAGs based on the quality standard settings (Table S3). Next, the high-quality genomes were clustered and compared based on genome-wide nucleotide diversity, and the best representative genome was selected for each cluster. A total of 589 SGBs were extracted, reaching an average mappability of 78.56 ± 6.46% of metagenomic reads per sample (Figure 1a). The high mappability suggested that the majority of genomic content in the samples’ microbiome corresponded to already known gut microbial community.

**Figure 1.**
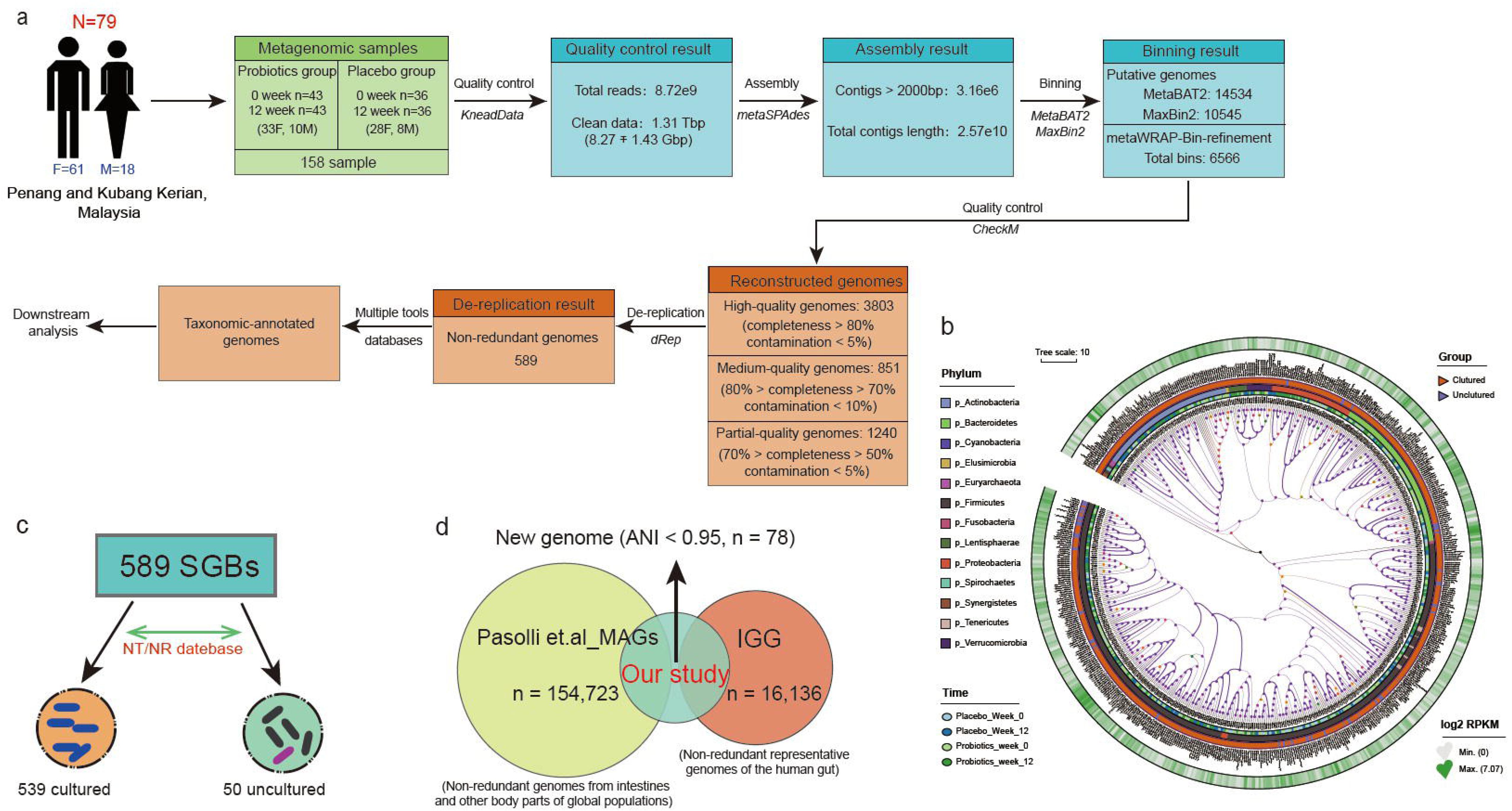
Metagenomic reconstruction pipeline and genomic characteristics. (a) The stepwise pipeline of metagenomic assembly, binning, reconstruction, and de-replication resulted in a total of 589 high-quality species-level genome bins (SGBs). (b) Phylogenetic placement of the 589 high-quality SGBs. The heatmap of the tree shows the log2-fold change in the RPKM of the SGBs during the trial. (c) Among the 589 SGBs, 539 and 50 were previously cultured and uncultured, respectively. (d) The novelty of the dataset and the 78 SGBs were confirmed by genomic comparison across the IGG database (a collection of 16,136 non-redundant representative human gut genomes) and the Pasolli et.al MAGs database (a dataset collection of 154,723 non-redundant genomes from gut and other body parts of the global population). These 78 SGBs were genomes reconstructed here and without overlapping with existing isolates or metagenomically assembled genomes of the compared human microbiome datasets.

The mapped SGBs were distributed across 13 phyla, 23 classes, 30 orders, 49 families, 91 genera, and 341 species. Most SGBs were assigned to the phylum Firmicutes (57.72%), followed by Bacteroidetes (14.60%), Actinobacteria (12.56%), and Proteobacteria (8.15%). Fifty SGBs remained unmapped and uncharacterized at the species level, and they represented the portion of uncultured species in our samples. These uncultured species mainly belonged to three phyla, i.e., Firmicutes (72%), Bacteroidetes (11.67%), and Proteobacteria (4%) (Figure 1b, 1c). To estimate the size of unexplored diversity in our SGBs dataset, the current dataset was cross-compared with the IGG dataset (https://github.com/snayfach/IGGdb; a collection of 16,136 non-redundant representative human gut genomes) and the Pasolli et.al MAGs dataset (a dataset collection of 154,723 non-redundant genomes from gut and other body parts of the global population). Seventy-eight SGBs did not match (shared < 95% average nucleotide similarity) any gut genome included in the two datasets, suggesting that they were novel species (Figure 1d; Table S4).

### Probiotic administration modulated the subjects’ gut microbiota composition

To assess the effect of probiotic consumption on the intestinal microbiome of stressed adults, an NMDS analysis (Bray-Curtis similarity index) was performed. Symbols representing samples at week 0 for both the placebo and P-8 groups were mainly distributed to the lower quadrants, while symbols representing samples at week 12 located mostly at the upper quadrants (Figure 2a). Such results suggested that obvious difference existed between the microbiota structure at weeks 0 and 12, which was supported by the result of ANOSIM. Further analysis by ANOSIM found no significant difference between the probiotic and placebo group at week 0 (R = 0.003, *p* = 0.354), but the two groups displayed significant difference at week 12 though with a relatively low R value (R = 0.041, *p* = 0.028). Interestingly, the placebo group showed a significantly larger difference in the Bray-Curtis distance between weeks 0 and 12 compared with the probiotic group (*P* < 0.001; Figure 2b). Moreover, comparing with week 0, the Shannon diversity index of the placebo group decreased significantly at week 12 (t-test; *p* < 0.05), but such decrease was non-significant for the probiotic group (Figure 2c). These results together suggested that although both the P-8 and placebo groups exhibited changes in alpha and beta diversity after the 12-week trial period, the probiotic-treatment resulted in smaller changes in the intestinal microbial diversity and microbiota structure.

**Figure 2.**
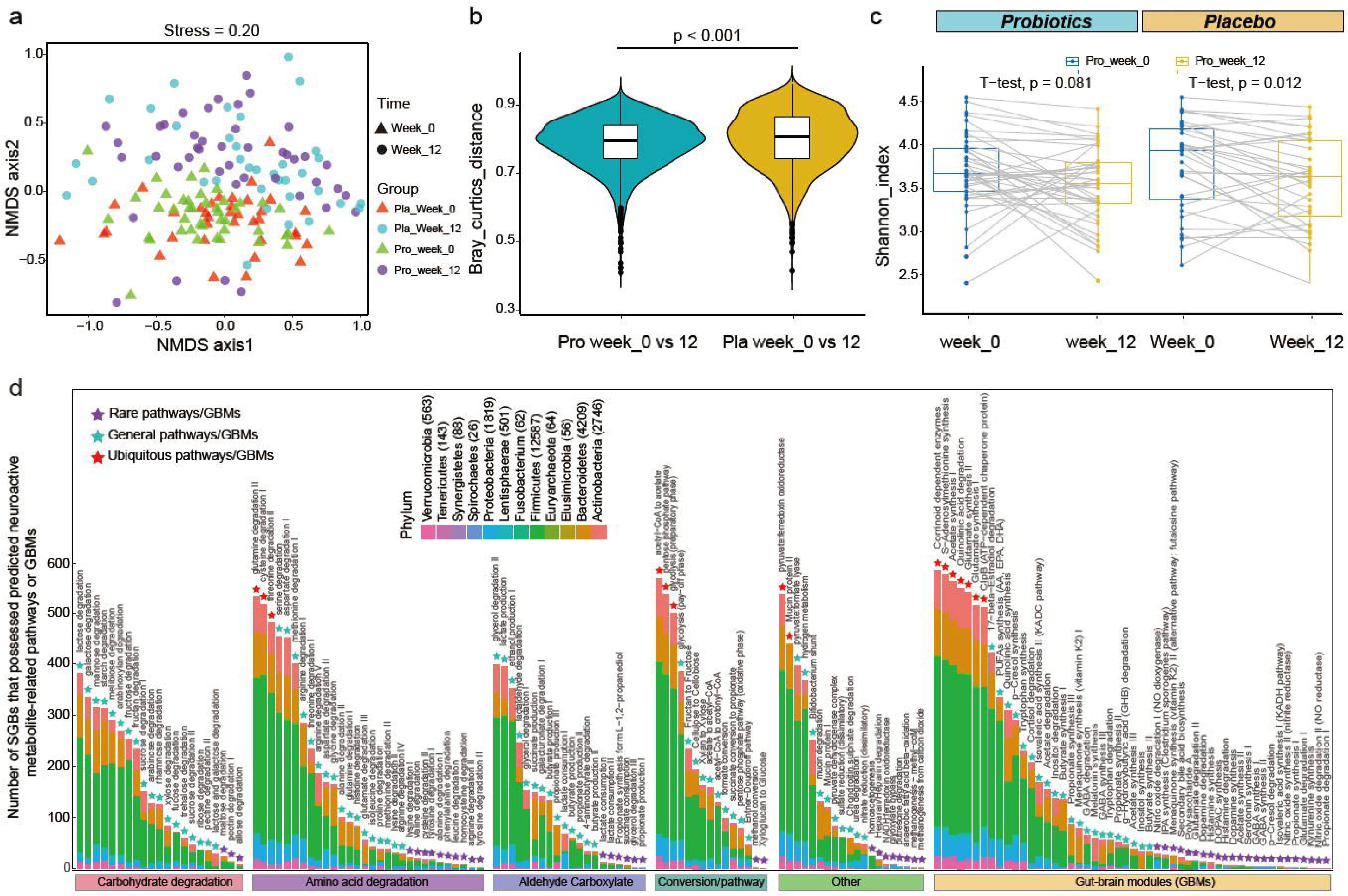
Structure, diversity, and functional features of intestinal microbiota in stressed adult. (a) Non-metric multidimensional scaling (NMDS) analysis (Bray-Curtis similarity index). Symbols representing samples of the placebo (Pla) and probiotic (Pro) groups are shown in different colors. Triangles and circles represent weeks 0 and 12 samples, respectively. (b) Significantly larger Bray-Curtis dissimilarity was observed in the gut microbiota (week 0 versus week 12) of the placebo group than the probiotic group. (c) Shannon diversity index of the two groups at two different time points. (d) Distribution of species-level genome bins (SGBs) that possessed predicted neuroactive metabolite-related pathways or gut brain modules (GBMs) across different phyla. The number written next to the phylum name indicates the quantity of SGBs distributed in that phylum. The purple, light blue and red triangle represent rare, general and ubiquitous pathways/GBMs, respectively.

To explore the probiotic-induced effect in the intestinal microbiota composition at a species-level, changes in the composition of SGBs of the placebo and probiotic groups were analyzed. The “core microbiota” comprised stable and permanent key microorganisms of the human intestine, and it was closely related to human health[43]. This study applied the concept of “core microbiota” to identify the “core SGBs” in the current dataset. Core SGBs were defined as SGBs present in more than 80% of the samples at week 0 and 12. A total of 13 core SGBs were identified, of which S24A.M011(*Fecalibacterium prausnitzii*) and S14A.M004 (*Fecalibacterium prausnitzii*), P08.M042 (*Agathobaculum butyriciproducens*), S37B.M011 (*uncultured Blautia sp.*), S25A.M021 (*Ruminococcus sp.* CAG:9-related_41_34), and H2_119.37 (*Bacteroides vulgatus*) showed no significant difference in abundance between the probiotic and placebo groups (Table S5). Six of the core SGBs responded to the probiotic treatment. At week 12, the P-8 group had significantly more S33A.M036 (*B. adolescence*), S4A.M008 (*B. longum*), H1_08.M012 (*F. prausnitzii*), and S34B.M002 (*Subdoligranulum sp*. 60_17) (*p* < 0.05), while significantly less H2_09.M016 (*Roseburia faecis*) and S22A.M013 (*Fusicatenibacter saccharivorans*; *p* < 0.05) were detected (Figure S1a). Then, ten SGBs (*B. longum*, *Megamonas funiformis*, *Subdoligranulum sp.* 60_17, and *Bacteroides sp*. et.al; including three uncultured species) were identified that had no significant difference between the placebo and the probiotic groups at week 0 but changed significantly at week 12. Differential abundant SGBs were also identified by comparing the abundance of all SGBs in the probiotic group before (week 0) and after (week 12) probiotic treatment, At week 12, ten SGBs (*Slackia isoflavoniconvertens*, *F. prausnitzii*, and *B. longum* et.al) were significantly enriched, whereas thirteen SGBs (*R. faecis*, uncultured *Ruminococcus sp.*, and *Dorea longicatena*) were depleted after the probiotic treatment (Table S5).

### Probiotic administration modulated gut microbiota-encoded neuroactivity-related pathways

To evaluate the specific effect of ingesting probiotics on the neuroactivity-related pathways of the intestinal microbiota, a genome-centric metabolic reconstruction approach was employed to identify changes in the pathways of the 589 identified SGBs using the MetaCyc and KEGG databases. The framework focused mainly on the human intestinal microbial metabolic pathways relating to the sugar fermentation, amino acid utilization, short-chain fatty acid synthesis (SCFAs) and degradation, gut enzyme conversion, and metabolism of human neuroactive compounds. These pathways of the current dataset belonged to 12 phyla, and most modules were distributed to Firmicutes (53.47%), Bacteroidetes (18.79%), and Actinobacteria (12.20%; Figure 2d). The two most ubiquitous pathways (prevalence > 80% among the SGBs) were acetyl-CoA to acetate (95.93%) and glutamine degradation II (89.98%). For gut-brain modules (GBMs), corrinoid dependent enzymes (96.69%) was most ubiquitous GMB. Some prevalent GBMs were detected in less than 5% of all SGBs, such as propionate degradation I, nitric oxide degradation II (NO reductase), and kynurenine synthesis; and these low occurring modules were mainly found in Proteobacteria and Actinobacteria (Table S6).

Thirty-two synthesis-related (Figure 3a) and seventeen degradation-related modules (Figure 3b) were found to be differential abundant between the probiotic and placebo groups at week 12. GBMs related to synthesis and degradation could be classified into seven subgroups according to their function characteristics. Significant differences were found between the two groups in the total abundance of SGBs encoding synthesis-related GBMs (SGBMs) at week 12, but not week 0 (*p* < 0.001). Comparing with the placebo group, the total abundance of SGBs involved in SGBMs V (organic acid synthesis) was significant higher in probiotic group at week 0 (*p* < 0.001), whereas the abundance of SGBs encoding SGBMs II (amino acid synthesis) and SGBMs III (ester synthesis) varied greatly between the P-8 and placebo groups at week 12 (*p* < 0.01; Figure S1b). Additionally, the abundance of SGBs involved in SGBMs IV (neurotransmitter II synthesis; contained three synthetic pathways of gamma-aminobutyric acid (GABA)) was higher in the probiotic group at week 12. The total abundance of SGBs possessing degradation-related GMBs (DGBMs) were not significantly different between weeks 0 and 12 in the probiotic and placebo groups. The abundance of SGBs encoding DGBMs I (neurotransmitter I degradation) and DGBMs IV (ester degradation) were not significantly different in the probiotic and placebo groups at week 0, but were significantly different at week 12, while other subgroups, e.g., DGBMs IV (neurotransmitter II degradation) and DGBMs VI (vitamins degradation), were changed little during the trial (Table S7).

**Figure 3.**
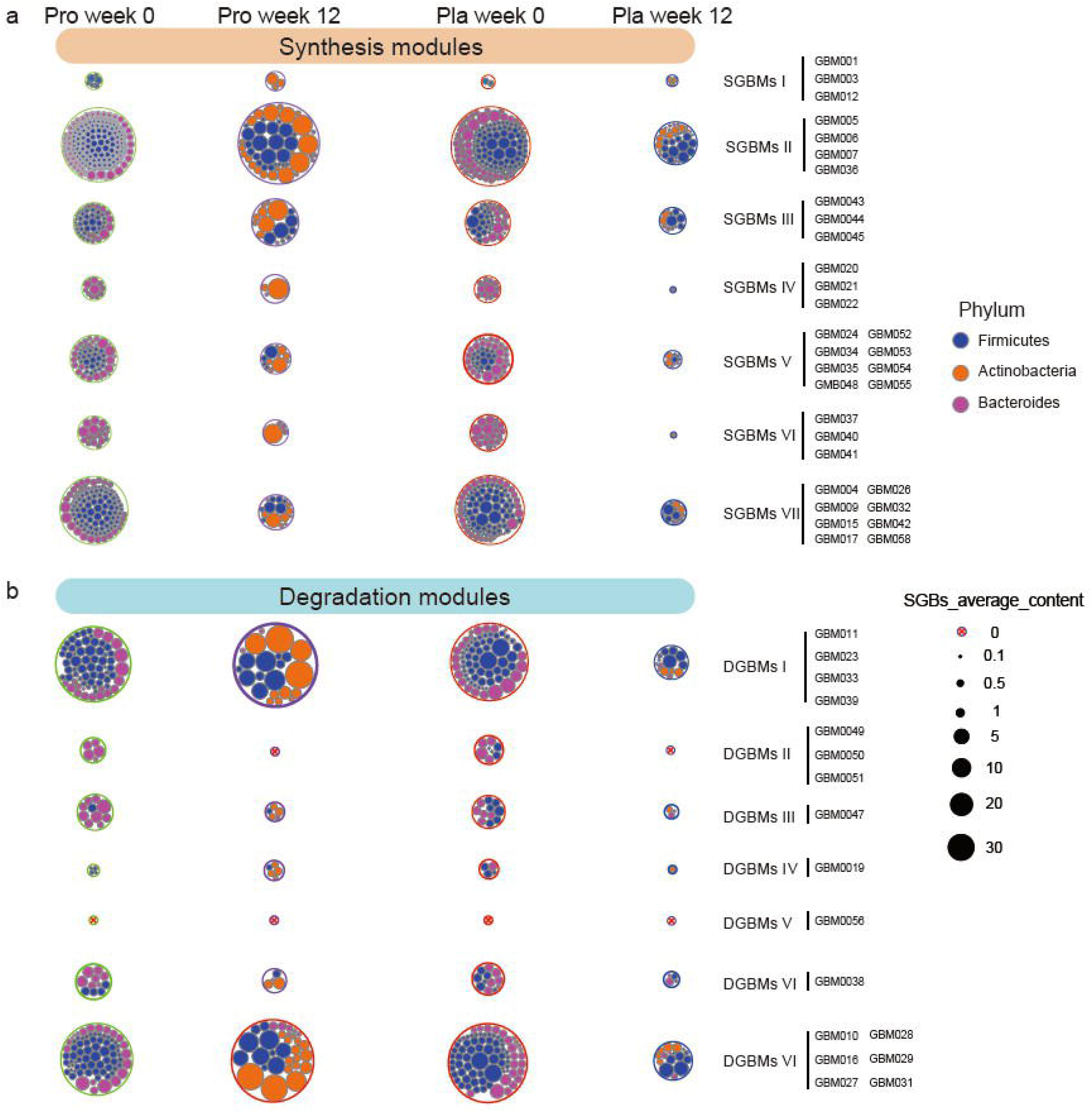
Distribution of gut brain modules (GBMs) at the phylum level. (a) The prevalence of different types of synthesis-related GBMs (SGBMs) or (b) degradation-related GBMs (DGBMs) and their compositions at the phylum level in the probiotic (pro) and placebo (pla) groups is shown. Identified SGBMs/DGBMs were classified into seven groups (I to VII) based on their predicted function of metabolite synthesis, respectively, and their associated species-level genome bins (SGBs) were assigned to corresponding phyla. The size of the outer circle represents the average abundance of corresponding SGBs.

### Probiotic administration modulated gut microbiota-related potential neuroactive compounds

Valles-Colomer et al[19]. developed a module-based analytical framework that facilitated targeted profiling of the microbial pathways involved in neuro-microbiome mediator metabolism. To profile the key potential neuroactive compounds, GBMs encoded by specific significant differential SGBs between the probiotic and placebo groups were analyzed by applying this framework (Figure S2a). At week 12, a higher diversity of SGB that were predicted to participate in menaquinone synthesis (vitamin K2) I synthesis, GABA and SCFAs (e.g. acetate, isovaleric acid) synthesis and consumption was found in the probiotic group, while a higher diversity of SGB was identified in the placebo group to participate in inositol degradation. Moreover, comparing with the placebo group, the subjects of P-8 group had more cortisol-degrading SGBs at week 12. Interestingly, more probiotic-receivers had histamine-synthesizing SGBs at week 12 (8 and 19 subjects at weeks 0 and 12, respectively; Tables S8)

### Microbiota neurometabolites features of the probiotic and placebo groups at two time points

To further establish the links between SGBs and potential neurometabolites, a metabolomic profiling prediction analysis was performed using MelonnPan. A total of 80 metabolites were predicted, among which deoxycholic acid, glutamate, and cholate were the three most abundant metabolites. The pattern of predicted fecal metabolomes of P-8 and placebo groups were analyzed by NMDS analysis to evaluate the effect of probiotic consumption on subjects’ intestinal metabolites. Although some overlapping occurred on the NMDS score plot, samples collected at different time points formed clear clustering pattern, with most of the symbols representing weeks 0 and 12 located at the right and left quadrants, respectively (stress = 0.08; Figure 4a). The Procrustes analysis showed a positive cooperativity between microbiome and metabolomic profiles (correlation = 0.386; *p* = 0.001; Figure 4b), suggesting a significant correlation between the abundances of SBGs and predicted metabolites. Furthermore, the ANOSIM found that significant changes occurred in the intestinal metabolite profiles after the 12-week trial period for both the P8 and placebo groups (*p* < 0.001; comparing between datasets at weeks 0 and 12 in both cases), but the metabolite profiles of the two groups were not significantly different at week 0. Specifically, 41 and 12 differential abundant metabolites were identified in the P-8 and placebo groups, respectively. These differential abundant metabolites showed significant changes in their predicted levels of abundance at week 12 compared with week 0. Some metabolites shared the same trend of change at the end of the clinical trial. For example, at week 12, both groups had significantly lower predicted levels of pantothenate, nicotinate, and lithocholate, while the predicted level of cytosine increased significantly after the 12-week period (*p* < 0.05 in all cases, t-test; Table S9). These changes were likely non-specific to the probiotic treatment. On the other hand, although there was no significant difference in the overall metabolite profile between the probiotic and placebo groups at week 12 based on NMDS and ANOSIM, the average predicted contents of cholate, arachidonic acid, creatine, threo-sphingosine, erythronic acid, and C18: 0 sphingomyelin were significantly higher in probiotic group (*p* < 0.05, t-test; Figure 4c).

**Figure 4.**
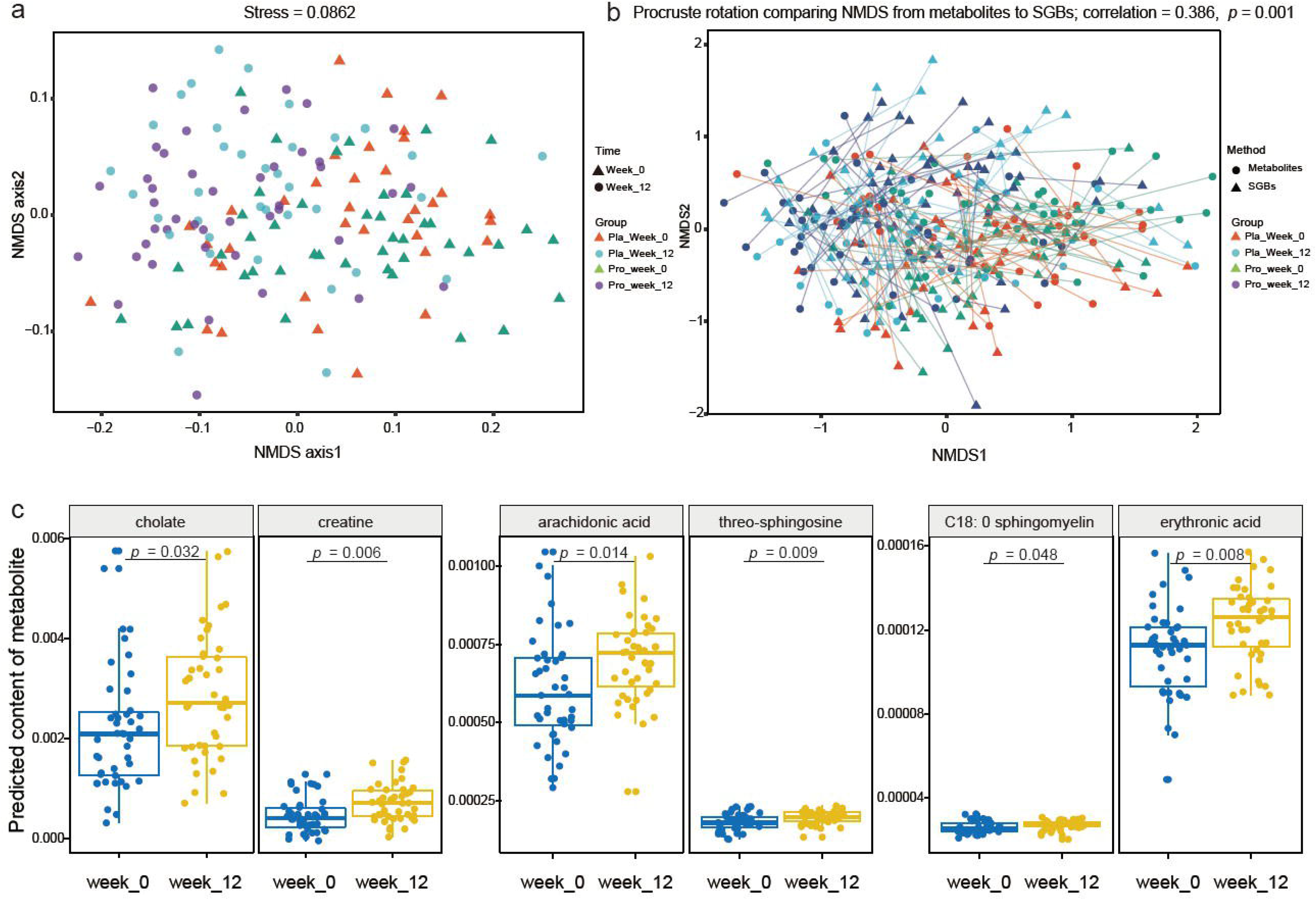
Differences in the predicted intestinal metabolomes of the probiotic (pro) and placebo (pla) groups at two time points. (a) Non-metric multidimensional scaling (NMDS, Bray-Curtis similarity index) and (b) Procrustes analyses were performed on the predicted microbiomes and metabolomes of the two groups of subjects at week 0 and week 12, showing a positive cooperativity between microbiome and metabolomic profiles (correlation = 0.386; p = 0.001). (c) Predicted differential neuroactive metabolites between week 0 and week 12 of the probiotic group.

## Discussion

The intestinal microbiota plays an indispensable role in human health, and it has recently become a biotherapeutic target for some common chronic diseases, such as metabolic syndromes, diabetes, obesity, and neurodegenerative diseases. Mounting evidence has shown that probiotic therapy creates a healthy intestinal environment by balancing bacterial populations and promoting favorable metabolism, which further highlights the role of bidirectional communication between the gut-brain axis[12, 44]. However, most previous studies have been performed on animal models (such as rodents, swine), and the human clinical data were not as convincing as animal models[45]. Furthermore, the lack of corresponding human intestinal reference genomes hinders a thorough understanding of the function and neuroactive potential of the intestinal microbiota. Our previous animal and human trials have shown beneficial effects of consuming *L. plantarum* P8 in regulating the colonic environment and gut microbiota, and ameliorating stress and anxiety symptoms in stressed population[25, 46]. However, the mechanisms behind the beneficial effects remained unclear. Thus, the current study was performed as a follow-up work to investigate the mechanism of relief of stress and anxiety symptoms from the perspectives of gut microbiota and neuroactive potential.

It is well known that psychological stress may cause bowel dysfunction and increase in gut permeability, which would in turn affect the gut microbiota community structure and activity[47]. The present research found that the intestinal microbiota structure of subjects under moderate level of stress changed during the trial, accompanied by a significant decrease in gut microbiota biodiversity and compositional changes in some species. Similar results have been reported in previous animal studies, mild and chronic stress shifted the caecum and fecal microbiota composition[48]. Moreover, even short-term exposure to stress could alter the gut microbiota profile at the family level[49]. Thus, the observation of stress-induced changes in microbiota composition and decreases in gut microbial diversity is consistent between different human and animal studies[50]. Several works showed that modulating the intestinal microbiota through prebiotic and/or probiotic interventions could alleviate stress-related behaviors, including anxiety and depression symptoms[12, 24]. Similar therapeutic effects were seen in the current study by P-8 intervention. It was interesting to note that the gut microbial diversity was smaller in the placebo group compared with the P-8 group after the 12-week trial. Another double-blind, placebo-controlled clinical trial of probiotics also found that the relief of stress-related psychological, physiological, and physical discomforts was accompanied by an increase in the gut microbiota diversity of volunteers after 8-week intervention[51]. Generally, a high diversity of gut microbiota has been recognized to be important for maintaining a healthy physiological state. Unhealthy conditions like gastrointestinal discomforts, Crohn’s disease, and some cancers have been associated with obvious reduction in gut microbial diversity; and gut dysbiosis could even partially perturb the normal function of energy metabolism, immunity, and oxidative stress, thus altering the health and performance of athletes[52]. Shade (2017) also argued against the assumption that a high diversity is implicitly a good or desirable outcome for the microbial communities or its relationships with emergent properties of a community, e.g., stability and productivity[53].

On the other hand, at the end of the 12-week probiotics treatment, the relative abundances of some SGBs increased significantly (e.g., *B. adolescent*, *B. longum*, and *F. prausnitzii*), while others decreased significantly (e.g., *R. faecis* and *Fusicatenibacter saccharivorans*). Some bacterial genera have been found to correlate positively with brain health, e.g., *Lactobacillus*, *Bifidobacterium*, and *Faecalibacterium*; some species/strains have shown regulatory and protective effects on neurological diseases[54]. For example, *B. adolescence* has been found to exert an anxiolytic and antidepressant effect by rebalancing the gut microbiota and reducing inflammatory cytokines[55]. Intake of *B. longum* 1714 not only ameliorated the physiological and psychological responses to an acute stressor, but also improved the visuospatial memory performance in healthy human adults[16]. Additionally, the administration of *F. prausnitzii* exerted preventive and therapeutic effects on depression-like and anxiety-like behavior in rats[56]. Our study also found negatively correlations between *B. adolescent*, *B. longum*, and *F. prausnitzii* with the symptom scores of stress and anxiety, while *R. faecis* correlated positively with these scores (Figure S2b). These results suggested that a diverse gut microbiota alone might not be enough to confer health-promoting effects to the host, but the presence of specific (groups of) functional gut microbes would be as important.

In recent years, multiple direct and indirect pathways that facilitate the communication between the gut and the brain have been identified, which involve the interactions between the vagus nerve, the immune system, the endocrine system, and microbial originated metabolites and neurotransmitters[54]. The gut microbial communities have been shown to produce and/or consume various intestinal neuroactive compounds, including dopamine, serotonin, histamine, SCFAs, GABA, and various neurohormones[8]. However, functional interpretation of metagenomic data in the microbiota-gut-brain context is still challenging, thus, this study investigated probiotic-specific modulation of the neuro-microbiome mediator metabolism by two different platforms, namely the module-based analytical framework developed by Valles-Colomer et al.[19] and MelonnPan[42]. Our data showed that the 12-week P-8 treatment resulted in significant increases in the diversity of SGBs that were involved in pathways associated with neuroactive potential, namely, menaquinone synthesis (vitamin K2) I, SCFAs and GABA metabolism. Vitamin K2 is an essential lipid-soluble vitamin that plays an important role in blood clotting and anti-inflammatory, which can be synthesized by a variety of bacteria in the human intestine. There is sufficient evidence that vitamin K2 has a protective effect on the nervous system and can be used to reduce the severity of relapses in patients with multiple sclerosis[57]. In addition, it has been found to prevent mitochondrial damage in individuals with drosophila, a pathology associated with Parkinson’s disease[58]. SCFAs (mainly including acetate, propionate and butyrate) are the products of intestinal bacterial fermentation and play a key role in neuroimmune endocrine regulation. Studies have found that acetate can alter the levels of the neurotransmitter glutamate, glutamine and GABA in the hypothalamus, and promote the development and maturation of brain glial cells[59]. In addition, a study shown that isovaleric acid, valeric acid, and acetate affect the pathogenesis of AD by disturbing the activations of microglia and astrocyte, help to reduce inflammation[60]. The *Fecalibacterium* genus is commonly recognized as a desirable gut SCFAs-producer, and it correlated with higher quality of life indicators[19], which is also consistent with our findings. GABA, is known to be produced in large amounts by intestinal bacteria; *Bifidobacterium* and *Lactobacillus* are common GABA-producers[8]. Our data showed that probiotic administration enriched the gut level of *B. adolescent* that encoded GABA synthesis pathways, and such change could be one reason for the improvement of depression and anxiety-related symptoms seen in the subjects in this study. Bravo et al. reported that ingesting *L. rhamnosus* reduced depression and anxiety behavior in mice in a vagus-dependent manner, as vagotomy prevented the anxiolytic and antidepressant effects of *L. rhamnosus* ingestion. Meanwhile, the symptom improvement was accompanied by an increase in GABAergic activity[61]. The most important neural pathway that communicates between the gut and the brain is the vagus nerve, and bacterial derived-neurotransmitters (e.g., GABA, histamine) transmit nerve inputs to the brain through the vagus ascending fibers[9]. Interestingly, microbial originated histamine could also transmit neural signal via the vagus nerve; however, how host histamine affects the physiology of the host remains to be uncovered[62]. Compared with the placebo group, a marginal decline was observed in plasma cortisol in the probiotic group, accompanied by enrichment of intestinal SGBs that involved in cortisol degradation. Cortisol is the primary stress hormone, which is connected to stress response like increasing anxiety and depression. It is known that some gut bacteria could reduce the level of cortisol, as evidenced by the decline in the urinary free cortisol level in subjects who took probiotics regularly[63].

Furthermore, neuroactive metabolites of the microbiota, such as bile acids and unsaturated fatty acids, also constitute a pathway of information flow between the gut and the brain[5]. Our data showed that the predicted levels of cholate, arachidonic acid, and C18:0 sphingomyelin increased significantly after probiotic treatment, these metabolites are known to be secreted by some gut bacteria. Bile acids help regulate the intestinal permeability and blood-brain barrier, and some secondary bile acids (such as cholate) can also stimulate 5-hydroxytryptamine (5-HT) biosynthesis[64]. Another study found that ingesting probiotics significantly increased the concentration of arachidonic acid in rodents. Arachidonic acid is a vital substance for the brain and optic nerve development, and it plays important roles in cognitive processes such as memory and learning[65].

The immune system plays an important role in the dynamic equilibrium of the gut-brain-axis. Peripheral administration of pro-inflammatory cytokines induced a variety of depression-like behaviors in rodents[66], while probiotics might exert immunomodulatory effects to counter stress response by generating T regulatory cell populations and secreting cytokines[67]. Our results found higher magnitudes of reduction in plasma pro-inflammatory cytokines, e.g. IFN-γ, TNF-α, and IL-1◻, in the probiotic group compared with the placebo group at week 12, which was consistent with the observation of published literature. Thus, the balance of intestinal microbiota may closely regulate host inflammatory response, activate intestinal and circulating immune pathways, subsequently influencing host mood and brain functions[9] (Figure 5).

**Figure 5.**
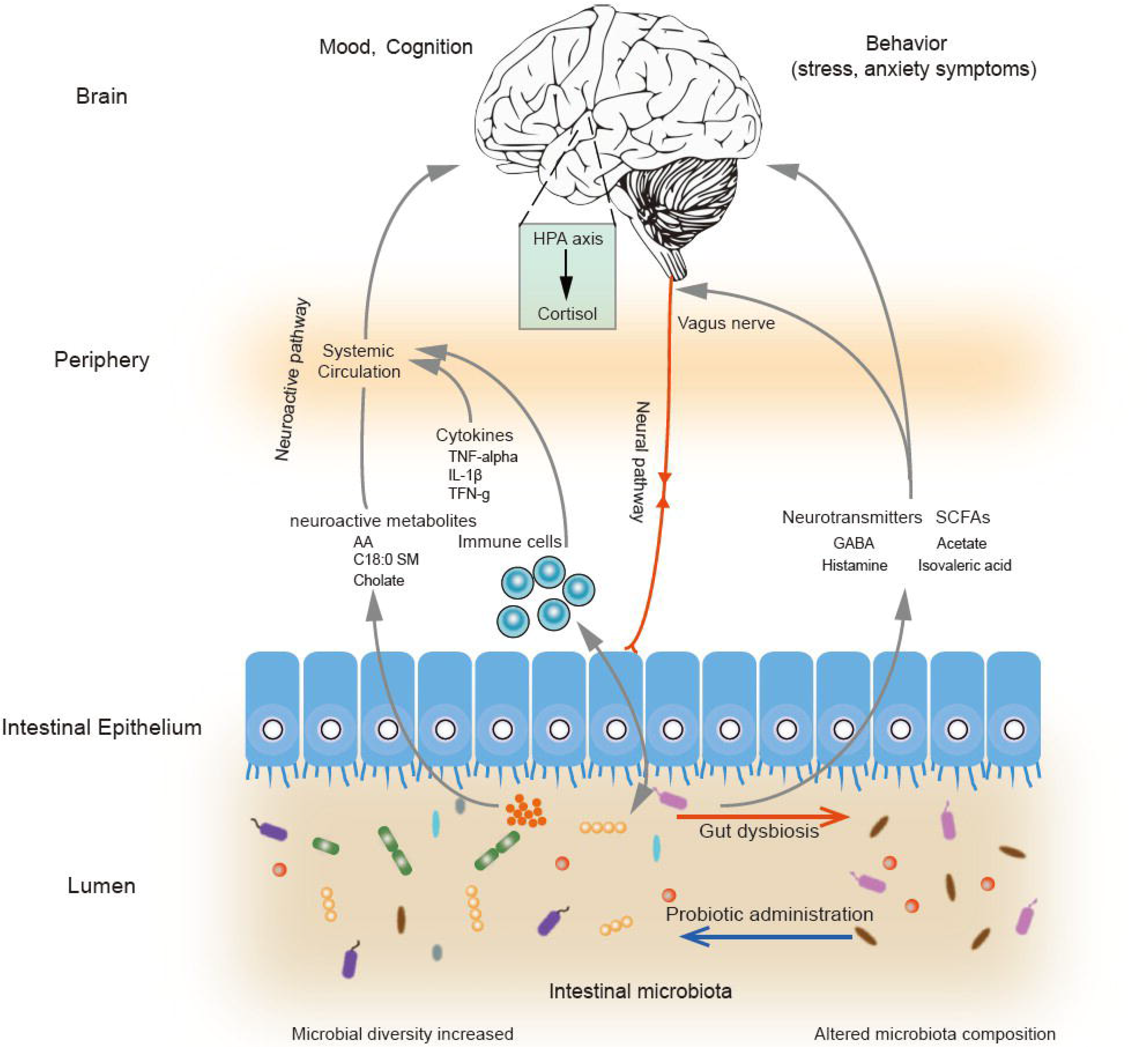
Schematic diagram illustrating key probiotic-driven pathways that modulated the gut-brain-axis and host response.

Collectively, P8 administration promoted the production and secretion of some key neurotransmitters and neuroactive metabolites in the intestine via regulating the gut microbiota. By such mechanisms, the intestinal microbiota regulated the gut-brain-axis directly via the vagus nerve and indirectly by means of cytokines and microbial originated metabolites. The current study revealed the potential mechanisms of probiotics in alleviating stress and anxiety symptoms in stressed adults. Our study supports that future development of psychobiotics targeting the intestinal microbiota for managing stress and anxiety symptoms would be valuable.

## Ethics approval and consent to participate

The study was reviewed and approved by the Ethics Committee of the Universiti Sains Malaysia, and informed consent was obtained from all subjects recruited from Penang and Kubang Kerian, Malaysia before they enrolled in the study.

## Authors’ contributions

MTL and HZ conceived and designed the experiments. JH and MT performed the study and analyzed the data. MT wrote the manuscript. LYK, ZJC, and SZH critically revised the manuscript. All authors read and approved the final manuscript.

## Figure Legends

**Figure S1.** Changes in core SGBs and synthesis-related gut-brain modules after probiotic treatment.

(a) Six of the core SGBs responded to the probiotic treatment. Compared with the placebo group, four core SGBs significantly increased and two decreased significantly after 12-week of probiotics treatment. (b) SGBs encoding synthesis-related gut-brain modules having significant different abundances between probiotic (Pro) and placebo (Pla) groups.

**Figure S2.** Changes of subjects’ neuroactive compounds and the correlation between differential SGBs and clinical scores under different treatment. (a) Gut-brain modules (GBMs) encoded by specific significant differential SGBs between the probiotic and the placebo groups. The circle, square represent enriched and depleted GBMs in the two groups at 12-week, respectively. (b) Correlation between differential SGBs and stress, anxiety, depression and total score in the probiotic and the placebo groups.

