## Supplementary figures and images for "Probiotic consumption relieved human stress and anxiety symptoms via modulating the gut microbiota and neuroactive potential"

### FigureS1.jpg

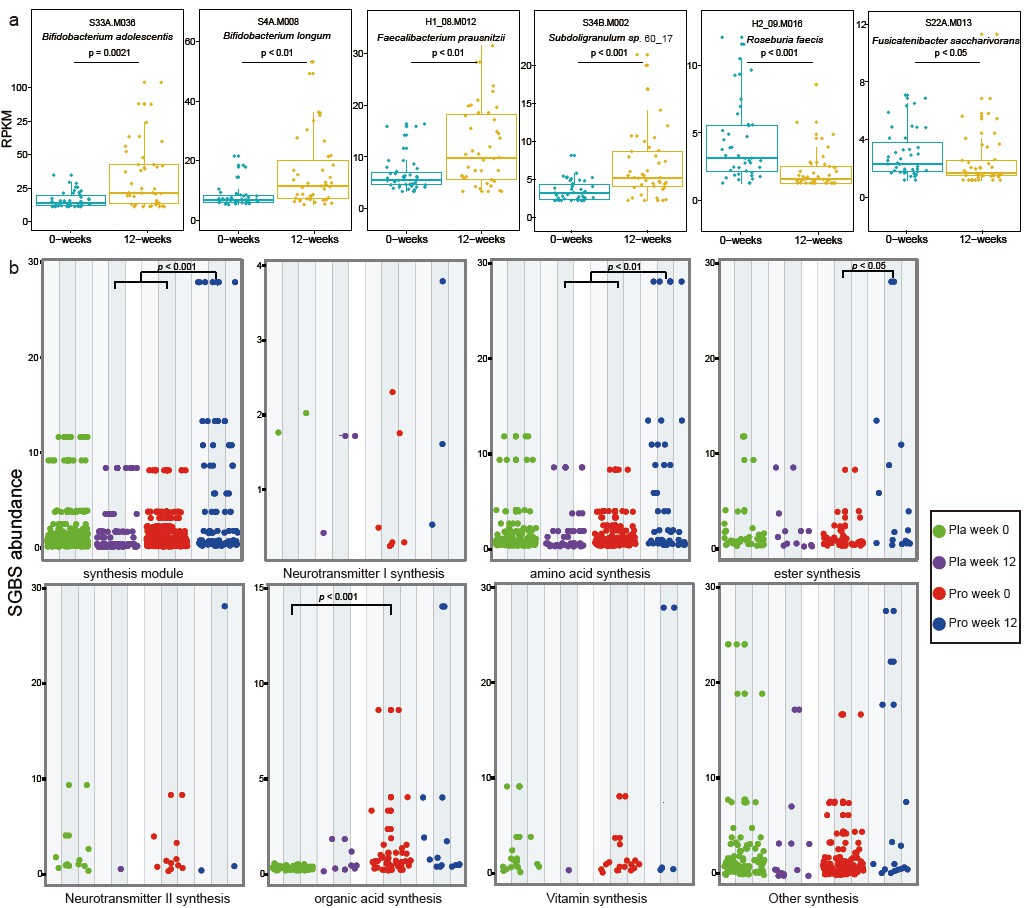

### FigureS2.jpg

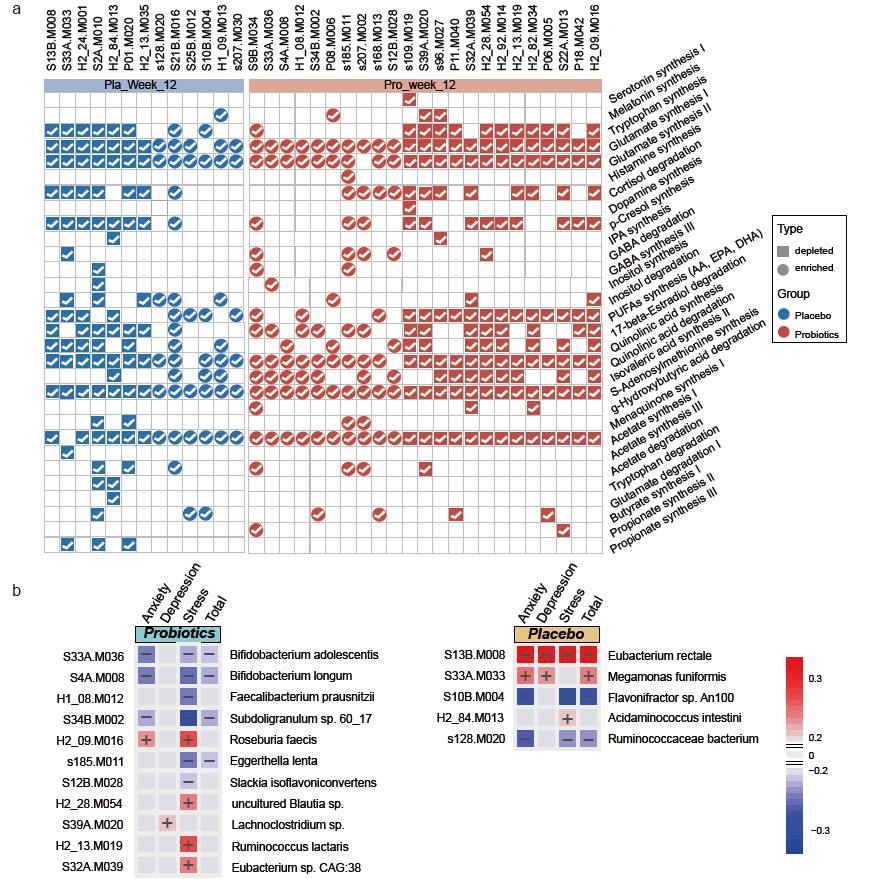
